# Taurine promotes axonal regeneration after a complete spinal cord injury in lampreys

**DOI:** 10.1101/655191

**Authors:** Daniel Sobrido-Cameán, Blanca Fernández-López, Natividad Pereiro, Anunciación Lafuente, María Celina Rodicio, Antón Barreiro-Iglesias

## Abstract

Taurine is one of the most abundant free amino acids in the brain. It is well known that taurine protects the brain from further damage after a traumatic event. However, only a few *ex vivo* studies have looked at the possible role of taurine in the regulation of axon regeneration after injury. Here, we aimed to reveal the possible role for taurine in the modulation of axonal regeneration following a complete spinal cord injury (SCI) using lampreys as an animal model. The brainstem of lampreys contains several individually identifiable descending neurons that differ greatly in their capacity for axonal regeneration after SCI. This offers a convenient model to promote or inhibit axonal regrowth in the same *in vivo* preparation. First, we carried out high performance liquid chromatography experiments to measure taurine levels in the spinal cord following SCI. Our results revealed a statistically significant increase in taurine levels 4 weeks post lesion, which suggested that taurine might have a positive effect on axonal regrowth. Based on these results, we decided to apply an acute taurine treatment at the site of injury to study its effect on axon regeneration. Results from these experiments show that an acute taurine treatment enhances axonal regeneration following SCI in lampreys. This offers a novel way to try to promote axon regeneration after nervous system injuries in mammalian models.

## Main text

In mammals, including humans, spinal cord injury (SCI) leads to permanent disability and an irreversible loss of sensorial, autonomic and motor functions below the lesion site. In contrast to mammals, lampreys recover locomotion spontaneously several weeks after a complete SCI.^1^ In spite of the amazing regenerative capacity of the central nervous system (CNS) of lampreys, not all neurons show the same abilities for axon regeneration. Among the brain descending neurons of lampreys there are 36 individually identifiable giant descending neurons. These identifiable descending neurons vary greatly in their regenerative abilities following a complete SCI,^2^ even when their axons run in similar paths in a spinal cord that is permissive for axonal regrowth. So, lampreys offer an interesting vertebrate model to study the molecular mechanisms that control axon regeneration.

Taurine is one of the most abundant free amino acids in the brain,^3^ and has various physiological functions, including osmoregulation and modulation of intracellular calcium levels.^4^ Many studies have also shown that taurine can protect the brain from traumatic damage.^5^ However, only a few studies have suggested that taurine could also promote axon regeneration after injury. For example, a couple of *ex vivo* studies in goldfish have shown that taurine can increase neurite growth in retinal explants with a prior crush injury to the optic nerve.^6, 7, 8, 9^ As far as we are aware, no studies have looked at the effects of taurine in axon regeneration *in vivo* in any vertebrate. Taurine concentrations are altered in the spinal cord after SCI in rats.^10, 11, 12^ Interestingly, methylprednisolone, which is the only pharmacological therapy used for the treatment of traumatic SCIs, ^13^ enhances axon regeneration after SCI,^14, 15^ and is known to cause an increase in the concentration of taurine in the spinal cord.^12^

Taurine is present in ependymal cells, radial glia and astrocytes of the lamprey spinal cord.^16^ Intracellular recordings of giant reticulospinal neurons of the sea lamprey have shown that these are hyperpolarized after taurine application.^17^ In vertebrates, taurine can act as an agonist of a variety of neurotransmitter receptors, including GABA receptors.^18^ Interestingly, recent work of our group has shown that endogenous GABA promotes axonal regeneration of identifiable reticulospinal neurons following SCI in lampreys.^19^ The beneficial effects of GABA appear to be mainly mediated through the activation of GABAB receptors in injured neurons;^19^ although, recent work has also shown that the activation of GABAA receptors by muscimol inhibits caspase activation in identifiable neurons after SCI in lampreys.^20^ Taken together, these data prompted us to study the possible effect of taurine in axon regeneration after SCI in lampreys. Here, we show that taurine levels increase 4 weeks after a complete SCI in the spinal cord of lampreys and that an acute taurine treatment promotes axon regeneration.

Mature and developmentally stable larval sea lampreys, *Petromyzon marinus* L. (n = 158; between 95 and 120 mm in body length, 5 to 7 years of age), were used in the present study. Animals were deeply anaesthetized with 0.1% MS-222 (Sigma, St. Louis, MO) in lamprey Ringer solution (137 mM NaCl, 2.9 mM KCl, 2.1 mM CaCl^2^, 2 mM HEPES; pH 7.4) before all experimental procedures and euthanized by decapitation at the end of the experiments. All experiments involving animals were approved by the Bioethics Committee at the University of Santiago de Compostela and the *Consellería do Medio Rural e do Mar* of the *Xunta de Galicia* (license reference JLPV/IId; Spain) and were performed in accordance to European Union and Spanish guidelines on animal care and experimentation. Complete spinal cord transections were performed as previously described.^21^ Briefly, the rostral spinal cord was exposed from the dorsal midline at the level of the 5^th^ gill by making a longitudinal incision with a scalpel (#11). A complete spinal cord transection was performed with Castroviejo scissors and the spinal cord cut ends were visualized under the stereomicroscope. The animals were allowed to recover in individual fresh water tanks at 19.5 ºC for 2, 4, 10 or 11 weeks post-lesion (wpl).

The high-performance liquid chromatography (HPLC) method was used to measure the concentration of taurine in the spinal cord of control and injured (2, 4 and 10 wpl) animals (n = 35 per group). These time points were chosen because axon retraction predominates in the first 2 weeks after a complete SCI, axon re-growth starts to predominate at 4 wpl,^22^ and at 10 wpl descending axons have already regenerated through the site of injury and animals show normal appearing locomotion. ^23^ Each HPLC sample contained a pool of 7 pieces (from 7 animals) of spinal cord from 1 mm rostral to the site of injury to 1 mm caudal to the site of injury. We used 5 samples for each of the 4 experimental groups (control and 2, 4 and 10 wpl animals). The samples were sonicated in acetic acid and stored at - 80ºC until their use for HPLC. Taurine was separated and analysed using HPLC with fluorescence detection after precolumn derivatization with O-phtalaldehyde (OPA), as previously described.^24^ An aliquot of 20 μl of the tissue supernatant containing homoserine as internal standard was neutralized with OPA reagent (4 mM OPA, 10% methanol, 2.56 mM 2-mercaptoethanol, in 1.6 M potassium borate buffer, pH 9.5) for 1 min. After this period, the reaction was stopped by adding acetic acid (0.5% v/v). Samples were immediately loaded through a Rheodyne (model 7125) injector system with a 50 μl loop sample to reach a C18 reverse-phase column (4.6 mm i.d. ×150 mm, Nucleosil 5, 100 A) eluted with a mobile phase consisting of 0.1 M sodium acetate buffer (pH 5.5) containing 35% methanol, at a flow rate of 1 mL/min and at a pressure of 140 bar. The column was subsequently washed with the same buffer containing 70% methanol and re-equilibrated with the elution buffer before use. The HPLC system consisted of a solvent delivery system coupled to a filter fluorometer (excitation 340 nm, emission 455 nm). Taurine content was calculated from the chromatographic peak areas using standard curves and the internal standard. The total protein content was measured in a NanoSpectrophotometer. Normality of the HPLC data was determined by the Kolmogorov-Smirnov normality test. All data passed the normality test. The results of control versus 2, 4 and 10 wpl groups were analysed by One-way ANOVA and Dunnett’s multiple comparisons test to compare each post-lesion group with the control group. Our results revealed a significant increase in the concentration of taurine in the spinal cord at 4 wpl (One-way ANOVA, *p =* 0.0017; Fig. 1). At 10 wpl (when animals recover normal appearing locomotion; ^23^) taurine levels decreased in the spinal cord and were not significantly different to control levels. Giving the good regenerative ability of lampreys, these results suggested that taurine could have a beneficial effect on recovery from SCI.

**Figure 1.**
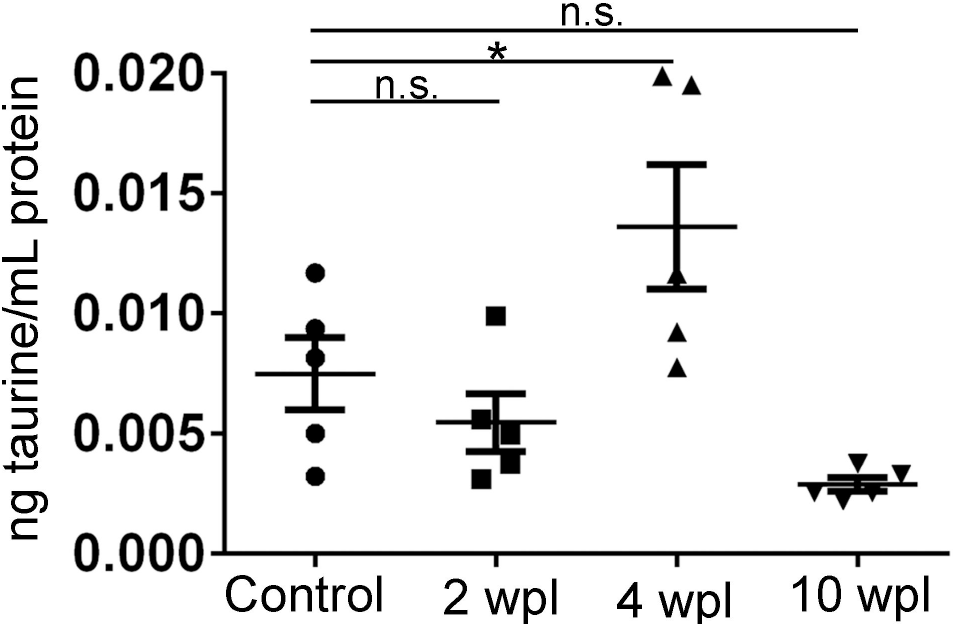
Graph showing taurine concentration in the spinal cord in control and injured animals (control: 0.0075 ± 0.01516 ng taurine/mL protein; 2wpl: 0.005468 ± 0.001195 ng taurine/mL protein; 4wpl: 0.01362 ± 0.002568 ng taurine/mL protein; 10wpl: 0.002876 ± 0.0002848 ng taurine/mL protein). Note the significant increase (asterisk) in taurine concentration at 4 wpl (Dunnett’s multiple comparisons test; *p* = 0.0419).

Based on the HPLC results, we decided to apply a taurine treatment at the time and site of the injury to see if supplying taurine acutely could further improve axon regeneration of sea lamprey descending neurons. Taurine was dissolved in lamprey Ringer solution at a concentration of 1 mM, soaked in a small piece of Gelfoam (Pfizer; New York, NY) and placed on top of the site of injury at the time of transection as previously described.^23^ Gelfoam soaked only in lamprey Ringer solution served as a control. Animals with a complete spinal cord transection were randomly assigned to either a vehicle treated control group or to taurine treated group. After the recovery period (11 wpl), a second complete spinal cord transection was performed 5 mm below the site of the original transection and the retrograde tracer Neurobiotin (NB, 322.8 Da molecular weight; Vector Labs, Burlingame, CA) was applied in the rostral end of the transected spinal cord with the aid of a minute pin (#000). The animals were allowed to recover at 19.5 ºC for 1 week to allow transport of the tracer from the application point to the neuronal soma of descending neurons (the M1, M2, M3, I1, I2, I3, I4, I5, I6, B1, B2, B3, B4, B5, B6, Mth and Mth’ neurons were analysed; see Figure 2A). Only neurons whose axons regenerate at least 5 mm below the site of injury are labelled by the tracer. Brains of these larvae were dissected out, and the posterior and cerebrotectal commissures of the brain were cut along the dorsal midline, and the alar plates were deflected laterally and pinned flat to a small strip of Sylgard (Dow Corning Co., USA). In some animals, the first portion of the spinal cord caudal to the obex was dissected out together with the brain to quantify the number of regenerated axons in the spinal cord. The brains/spinal cords were fixed with 4% PFA in TBS for 4 hours at 4 ºC. After washes in TBS, the brains/spinal cords were incubated at 4 ºC with Avidin D-FITC conjugated (Vector; Cat#: A-2001; 1:500) diluted in TBS containing 0.3% Triton X-100 for 2 days to reveal the presence of Neurobiotin. Brains were rinsed in TBS and distilled water and mounted with Mowiol. A spectral confocal microscope (model TCS-SP2; Leica, Wetzlar, Germany) was used to acquire images of the brains/spinal cords. The identify of regenerated (Neurobiotin labelled) identifiable descending neurons was determined for each brain. For axon quantification, we scanned the rostral spinal cord (starting at the obex) using a 20x objective. Then, we traced a horizontal line through the middle of the spinal cord confocal projection (see Fig. 2A) using the Fiji software,^25^ and manually quantified the number of axons crossing this line. Only the axons that regenerate through the site of injury (ascending or descending) are labelled with the tracer. Note that with this preparation we can only quantify the coarsest axons; therefore, it is likely that more axons actually regenerated both in control and treated animals. The experimenter was blinded during quantifications.

**Figure 2.**
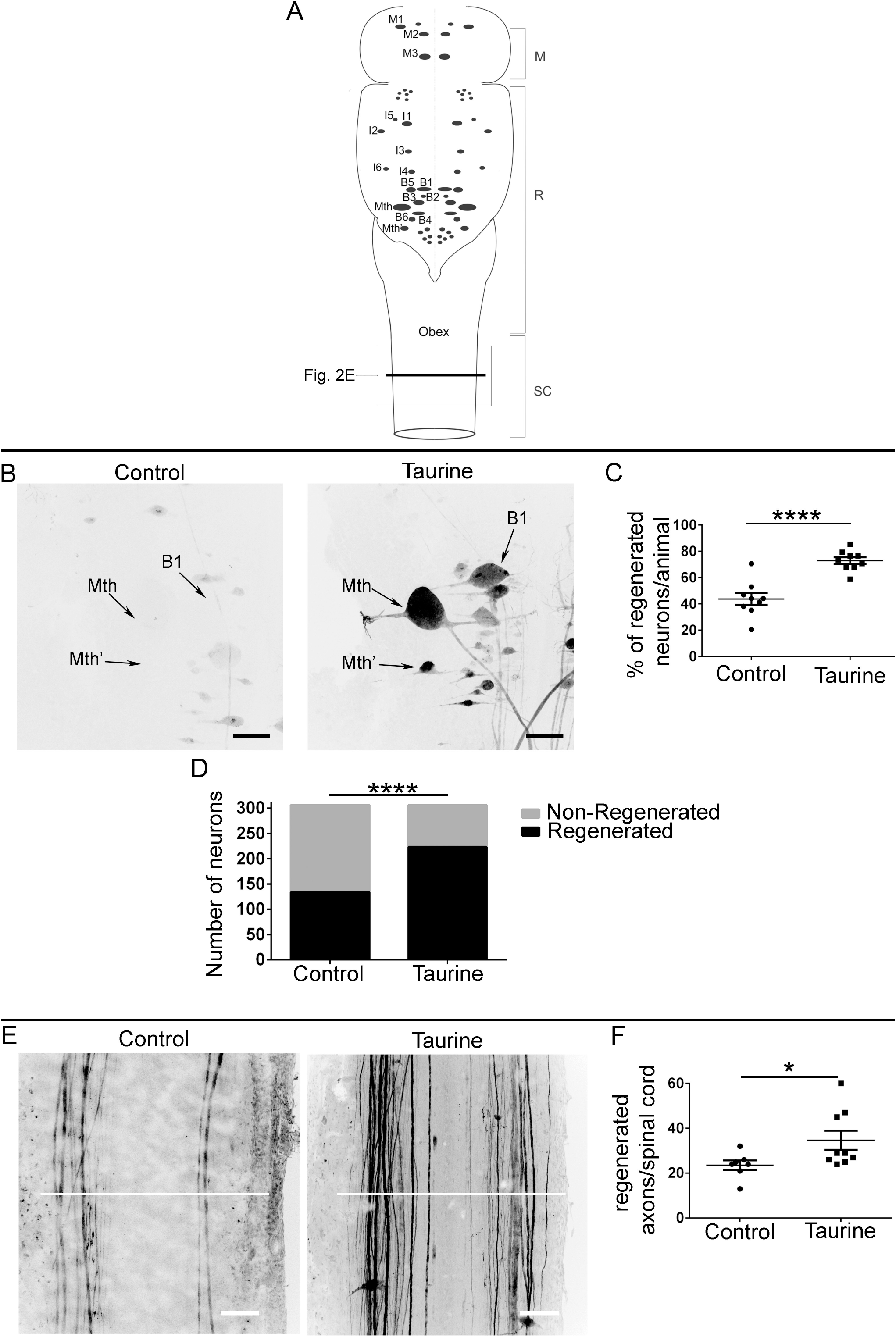
Taurine promotes axon regeneration after a complete SCI. **A**: Schematic drawing of a dorsal view of the sea lamprey brainstem showing the location of identifiable descending neurons (modified from ^13^). The spinal cord region shown in figure 2E is indicated with a square. The line used to count the number of regenerated axons in the spinal cord is also indicated. Abbreviations: M: mesencephalon; R: rhombencephalon; SC: spinal cord. **B**: Photomicrographs of whole-mounted brains showing regenerated identifiable neurons, as identified by retrograde tracer labelling, in control and taurine treated animals. Note the increased number of labelled (regenerated) identifiable neurons in taurine treated animals. **C**: Graph showing significant changes (asterisks) in the percentage of regenerated identifiable descending neurons per animal after the taurine treatment (control: 43.79 ± 4.505%; taurine: 72.88 ± 2.584%). **D**: Graph showing significant changes (asterisks) in the total number of regenerated neurons after the taurine treatment (control: 134 neurons regenerated, 172 non-regenerated; taurine: 223 neurons regenerated, 83 non-regenerated). **E**: Photomicrographs of whole-mounted spinal cords showing regenerated axons, as identified by tracer labelling, in control and taurine treated animals. Note the increased number of regenerated axons in taurine treated animals. The line used to count the number of regenerated axons in the spinal cord is also indicated. **F**: Graph showing significant changes (asterisk) in the number of regenerated axons per spinal cord after the taurine treatment (control: 23.57 ± 2.17 axons; taurine: 34.67 ± 4.259 axons). Rostral is up and scale bars: 100 µm in all photomicrographs.

At 11 wpl, the taurine treatment significantly improved axon regeneration of identifiable descending neurons of the sea lamprey (Fig. 2B-D); both when looking at the percentage of regenerated identifiable descending neurons per animal (Student t-test, *p* < 0.0001; Fig. 2B-C) or at the total number of regenerated neurons (Fisher’s exact test, *p* < 0,0001; Fig. 2B and D). Moreover, the acute taurine treatment significantly increased the number of regenerated axons in the spinal cord following a complete SCI (Mann Whitney test, *p* = 0.0289; Fig. 2E-F). These results show that supplying taurine acutely improves axon regeneration of sea lamprey neurons after SCI.

Our results show, for the first time, that taurine promotes axon regeneration following SCI. Importantly, this is shown at the level of individually identifiable neurons and in a model of complete SCI, which allows to study true axonal regeneration. Here, we showed that taurine concentration in the sea lamprey spinal cord is significantly increased at 4 wpl, which coincides with time in which axon re-growth starts to predominate.^22^ This observation prompted us to study whether taurine could be promoting axon regeneration following SCI. We confirmed the pro-regenerative effect of taurine by performing an acute taurine treatment at the time of SCI. This treatment resulted in a significant increase in axon regeneration at 11 wpl, which suggests that the acute taurine treatment could be suppressing or reducing axon retraction after axotomy and promoting axon re-growth. In future work, it would be interesting to decipher the molecular mechanisms behind the pro-regenerative effect of taurine in lampreys. Our results provide a possible new way to try to promote regeneration in mammalian pre-clinical models of SCI.

## Acknowledgments

Grant sponsors: Spanish Ministry of Economy and Competitiveness and the European Regional Development Fund 2007–2013 (Grant numbers: BFU2014-56300-P and BFU-2017-87079-P). A.B.-I. was supported by a grant from the crowdfunding platform *Precipita* (FECYT; Spanish Ministry of Economy and Competitiveness; grant number 2017-CP081). The authors would like to acknowledge the following individual donors of the crowdfunding campaign in *Precipita*: Blanca Fernández, Emilio Río, Guillermo Vivar, Pablo Pérez, Jorge Férnandez, Ignacio Valiño, Pago de los Centenarios, Eva Candal, María del Pilar Balsa, Jorge Faraldo, Isabel Rodríguez-Moldes, José Manuel López, Juan José Pita, María E. Cameán, Jesús Torres, José Pumares, Verónica Rodríguez, Sara López, Tania Villares Balsa, Rocío Lizcano, José García, Ana M. Cereijo, María Pardo, Nerea Santamaría, Carolina Hernández, Jesús López and María Maneiro. The authors also thank the staff of the Ximonde Biological Station for providing lampreys used in this study, and the Microscopy Service (University of Santiago de Compostela) and Dr. Mercedes Rivas Cascallar for confocal microscope facilities and help.

